# Machine Learning Identifies Novel Candidates for Drug Repurposing in Alzheimer’s Disease

**DOI:** 10.1101/2020.05.15.098749

**Authors:** Steve Rodriguez, Clemens Hug, Petar Todorov, Nienke Moret, Sarah A. Boswell, Kyle Evans, George Zhou, Nathan T. Johnson, Brad Hyman, Peter K. Sorger, Mark W. Albers, Artem Sokolov

## Abstract

Clinical trials of novel therapeutics for Alzheimer’s Disease (AD) have consumed a large amount of time and resources with largely negative results. Repurposing drugs already approved by the Food and Drug Administration (FDA) for another indication is a more rapid and less expensive option. Repurposing can yield a useful therapeutic and also accelerate proof of concept studies that ultimately lead to a new molecular entity. We present a novel machine learning framework, DRIAD (<u>D</u>rug <u>R</u>epurposing In <u>AD</u>), that quantifies potential associations between the pathology of AD severity (the Braak stage) and molecular mechanisms as encoded in lists of gene names. DRIAD was validated on gene lists known to be associated with AD from other studies and subsequently applied to evaluate lists of genes arising from perturbations in differentiated human neural cell cultures by 80 FDA-approved and clinically tested drugs, producing a ranked list of possible repurposing candidates. Top-scoring drugs were inspected for common trends among their nominal molecular targets and their “off-targets”, revealing a high prevalence of kinases from the Janus (JAK), Unc-51-like (ULK) and NIMA-related (NEK) families. These kinase families are known to modulate pathways related to innate immune signaling, autophagy, and microtubule formation and function, suggesting possible disease-modifying mechanisms of action. We propose that the DRIAD method can be used to nominate drugs that, after additional validation and identification of relevant pharmacodynamic biomarker(s), could be evaluated in a clinical trial.

## Introduction

Alzheimer’s Disease (AD) is a growing healthcare crisis with longer life expectancy as its principal risk factor. It is estimated that, in the absence of effective prevention and treatment options, disease prevalence will more than double over the next several decades: from 5.8 million individuals living with AD today in the US to a projected 13.8 million by 2050^1^. In addition to its direct impact on human health and welfare, the long-term care of affected individuals imposes a substantial economic burden^2^. Multiple efforts to develop disease-modifying therapeutics for AD, including 200 clinical trials to date, have been largely negative with many failures occurring due to excess toxicity and lack of efficacy^3^. Every failed clinical trial of a new molecular entity (NME) consumes substantial time and resources. In contrast, repurposing drugs already approved by the Food and Drug Administration (FDA) for another indication is less expensive and has a higher success rate (30%) as compared to development of an NME. The most obvious approach to repurposing is to use an existing drug in a new indication, perhaps at a different dose or in different formulation.^4^ An alternative is to use repurposing as a way of testing a therapeutic concept that could then be subsequently advanced via an NME. This is potentially valuable in the case of AD in which the underlying disease mechanisms remain poorly understood and the potential for multiple distinct disease etiologies exists. Repurposing drugs for AD has received increasing attention^5,6^, but approaches to date have been largely hypothesis-driven, based on overlap between an existing pharmacological mechanism of action (MOA) and a putative disease-causing mechanism^7^ or results of a clinical trial^8,9^. While some of these leads are promising, no successes have been reported to date.

As databases of drug information grow, informatics-based approaches to drug repurposing have emerged. Tools such as PREDICT^10^, Rephetio^11^, Connectivity Map^12^, and other methods^13,14^ seek to establish large-scale associations between perturbations induced by drugs and by disease processes, which can be mined for novel repurposing opportunities. For simplicity, we use the term *drug* in this work to broadly refer to both the FDA-approved chemical entities as well as clinically tested compounds and drug-like pre-clinical molecules (often referred to as “chemical probes”). One drawback of existing repurposing databases is that they are rarely disease specific and include data on drug mechanisms and disease pathways (typically transcriptional or proteomic signatures) obtained from diverse biological settings. This is potentially problematic in the case of a disease such as AD that is poorly understood and characterized by phenotypic^15^ and pathological heterogeneity^16^. We therefore sought to develop a repurposing approach that makes use of omic datasets on drug mechanisms and on disease, as obtained from individuals suffering from different stages of AD; such databases have been collected by the Accelerating Medicines Partnership - Alzheimer’s Disease (AMP-AD) effort^17^.

In this manuscript we describe DRIAD (Drug Repurposing In Alzheimer’s Disease), a novel machine learning framework that quantifies the association between the stage of AD (early, mid or late) as defined by Braak staging^18^ and any biological process or response that can be characterized by a list of gene names. Data characterizing AD were obtained from AMP-AD datasets and comprised mRNA expression profiles of postmortem brain specimens. Lists of gene names were obtained by using RNAseq to measure the responses of human neuronal cells to small molecule drugs and then identify differentially expressed genes (to generate *drug-associated gene lists*; DGLs). In the current work, we focus on kinase inhibitors because they are associated with strong transcriptional signatures and have relatively well annotated target spectra^19^. DRIAD uses a specific DGL for feature selection and then trains and evaluates a predictor of AD pathological stage from AMP-AD gene expression data. In this way, the relevance of a drug-induced perturbation of neurons (which is a reflection of drug mechanism of action) to the pathological processes underlying AD can be evaluated. Drugs whose DGLs resulted in the most accurate predictors were found to target proteins in signaling networks regulating innate immunity, autophagy, and microtubule dynamics; these represent novel pathways for a potential Alzheimer’s therapeutic^20^. The direction of this effect is not specified a priori and DRIAD will identify both disease-enhancing and disease-reducing drugs, a topic addressed in the discussion. We also show that DRIAD is agnostic to the type of disease, the length of the input gene list and its source, which can be a drug-induced perturbation, as in this work, the results of a previous study or manual annotation of biological mechanisms.

## Results

### Machine learning framework for identifying potential associations between gene lists and disease

When used for AD repurposing, DRIAD requires two types of inputs: (i) mRNA expression profiles from human brains at various stages of AD progression and (ii) a dataset comprising DGLs – lists of genes differentially expressed when neurons are exposed to a test panel of drugs. Human brain gene expression levels were taken from AMP-AD datasets^17^ provided by The Religious Orders Study and Memory and Aging Project (ROSMAP), The Mayo Clinic Brain Bank (MAYO) and The Mount Sinai/JJ Peters VA Medical Center Brain Bank (MSBB), each encompassing measurements from one or more regions of the brain (Fig. 1b). Braak staging scores^18^, assigned through neuropathological assessment, were used to group samples into three categories of disease progression: early (A; stages 1-2), intermediate (B; stages 3-4) and late (C; stages 5-6). This grouping recapitulates the spatio-temporal progression of neurofibrillary tangles from the entorhinal region to the hippocampus area and, subsequently, to neocortical association areas^21^.

**Fig. 1:**
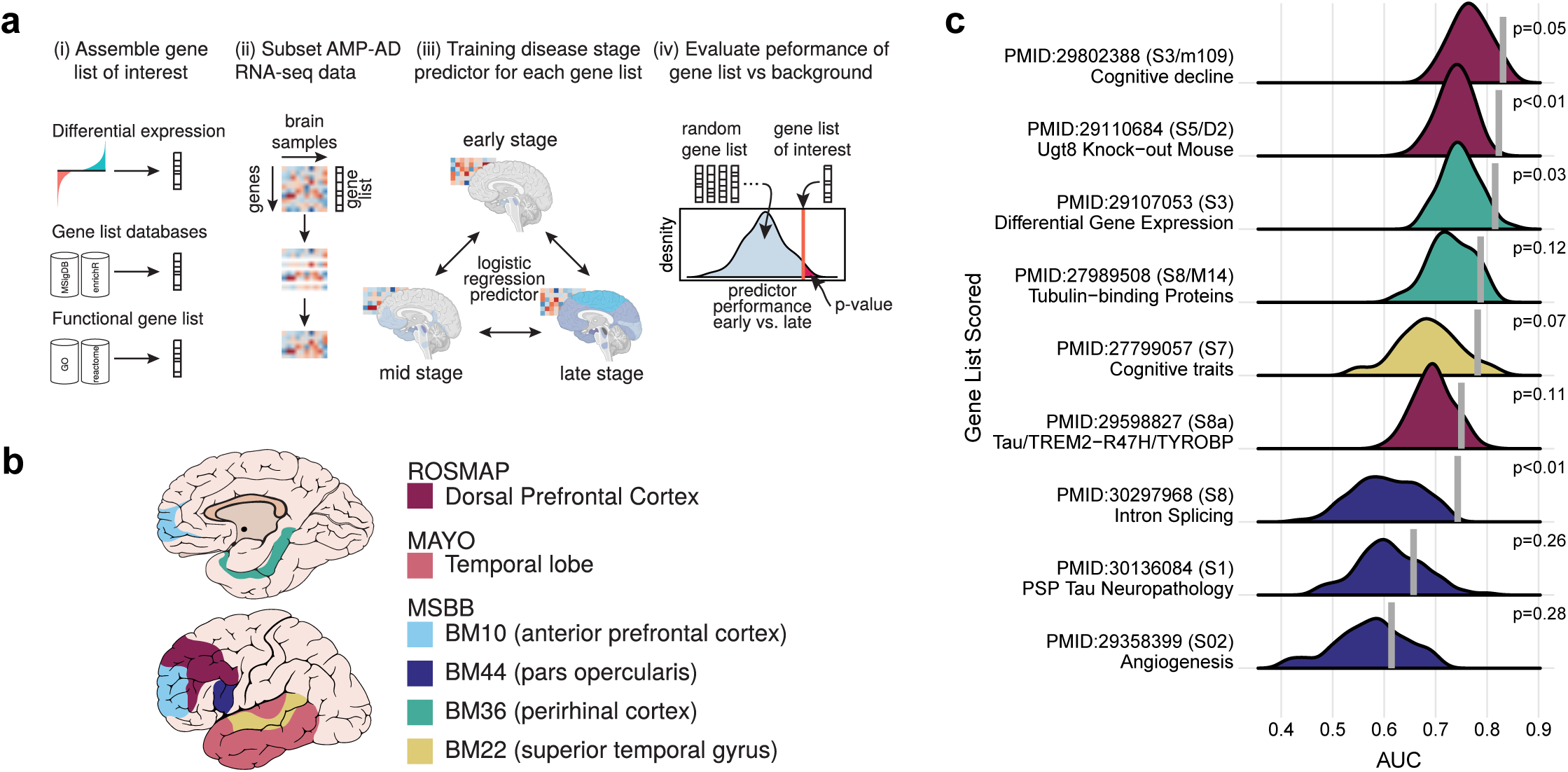
**a**, Overview of the machine learning framework used to establish potential associations between gene lists and Alzheimers Disease. (i) The framework accepts as input gene lists derived from experimental data or extracted from database resources or literature. (ii) Given a gene expression matrix, the framework subsamples it to a particular gene list of interest, and (iii) subsequently trains and evaluates through cross-validation a predictor of Braak stage of disease. (iv) The process is repeated for randomly-selected gene lists of equal lengths to determine whether predictor performance associated with the gene list of interest is significantly higher than whats expected by chance. **b**, AMP-AD datasets used by the machine learning framework. The three datasets used to evaluate the predictive power of gene lists are provided by The Religious Orders Study and Memory and Aging Project (ROSMAP), The Mayo Clinic Brain Bank (MAYO) and The Mount Sinai/JJ Peters VA Medical Center Brain Bank (MSBB). The schematic highlights regions of the brain that are represented in each dataset. The MSBB dataset spans four distinct regions, which are designated using Brodmann (BM) area codes. **c**, Performance of predictors trained on gene lists reported in previous studies of AMP-AD datasets. The predictors are evaluated for their ability to distinguish early-vs-late disease stages with performance reported as area under the ROC curve (AUC). The vertical line on each row denotes predictor performance associated with a gene list reported in the literature, while the background distribution is constructed over randomly-selected lists of matching lengths. Each row is annotated with the pubmed ID of the study, the supplemental resource that contained the gene list, and a short keyphrase providing functional context.

DRIAD trains and then evaluates a predictor that can recognize the A, B or C disease category from mRNA expression levels, restricting the predictor to those genes in the DGL (Fig. 1a) (See Methods). We consider it highly likely that the molecular processes involved in the initiation and progression of AD are obscured in end-stage RNAseq profiles as a result of the actions of myriad signaling pathways and feedback loops, leading to widespread transcriptional changes only indirectly associated with disease mechanism^22–25^. This is reflected in the fact that any randomly-selected list of genes in human brain gene expression profiles is weakly predictive of disease stage (Supplementary Fig. 1). We therefore sought lists of genes that outperformed random lists in predicting AD stage at a pre-specified level of significance. Statistical significance was assessed by repeatedly sampling the space of all gene names to create a background distribution of random gene lists (of the same length) against which to evaluate a DGL (Fig. 1a). An empirical p value was provided by the fraction of random gene lists that outperformed the DGL in the prediction task.

### Validation of DRIAD using gene lists associated with AD pathophysiology

To validate the DRIAD framework, lists of gene names previously reported in the literature to be associated with an aspect of AD progression^24,26–33^ were substitute for DGLs. Each input gene list was used to train DRIAD to distinguish between early (A) and late (C) stage disease based on mRNA levels in AMP-AD data. We found that most of the published lists of AD-associated genes outperformed randomly-selected lists of equal length (Fig. 1c). Thus, DRIAD effectively recapitulates previous attempts to identify genes and co-expression modules associated with disease severity. Similar results were obtained with human gene sets and mouse homologues previously reported to be associated with AD. For example, McKenzie, *et al*., used AMP-AD data to establish that *Ugt8* is a key regulator of oligodendrocyte function and reported a list of genes that are differentially expressed in the frontal cortex of a *Ugt8* knock-out mouse model^32^. DRIAD shows that these differentially expressed genes have a significant association with disease severity in AMP-AD data (Fig. 1c), providing further evidence that *Ugt8* may play a role in neurodegeneration in humans.

Similarly, Mostafavi, *et al*., identified a co-expression module of 390 genes that has a strong association with cognitive decline^26^. DRIAD confirms that a predictor trained to recognize pathological stage based on these 390 genes performs better than equivalent predictors trained on any arbitrary set of 390 genes chosen at random (Fig. 1c). In contrast, when we examined gene lists associated with Progressive Supranuclear Palsy (PSP) Tau Neuropathology and Angiogenesis, DRIAD found that they correlated with Braak staging no better random. This suggests that models trained using DRIAD can distinguish molecular features of tauopathies and pathologic processes not known to be relevant to AD. Consistency in performance was observed across datasets and brain regions, with one exception: whenever a predictor was trained on late-stage samples in the MAYO dataset, it substantially outperformed similar predictors evaluated on all other datasets (Fig. 2b, Supplementary Fig. 1). This suggests the presence of a strong batch effect in the late-stage MAYO samples that the predictors pick up in lieu of learning disease severity. Because the nature of this batch effect is unknown, we chose to exclude MAYO data in the current study. However, future studies may consider adjusting for batch effects of unknown origin using established methods from the literature^34–36^.

**Fig. 2:**
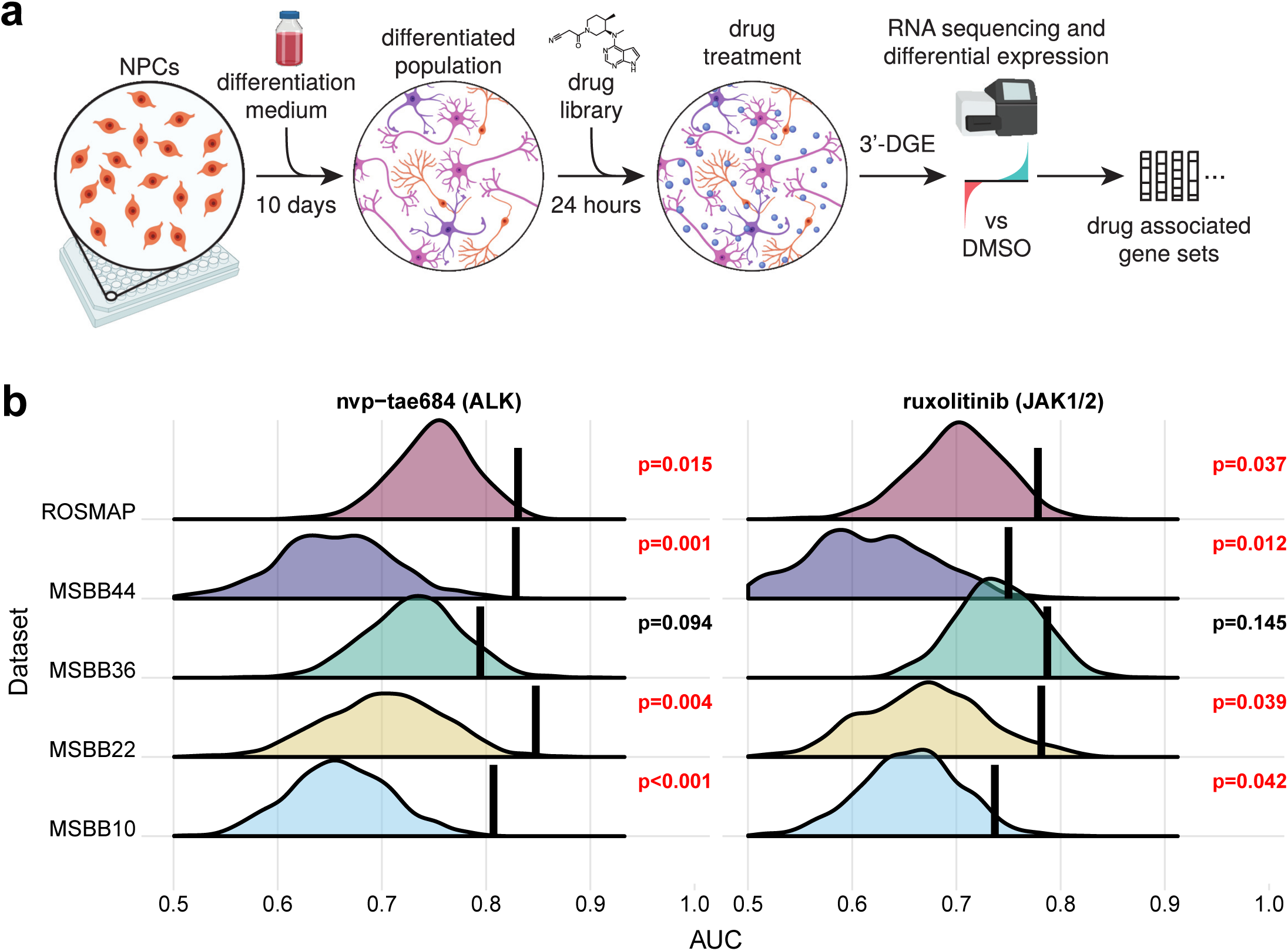
**a**, Overview of the 3’ DGE experimental protocol used to derive drug-associated gene expression signatures. ReNcell VM human neural progenitor cells were plated and differentiated for 10 days, resulting in a mixed cell population of neurons, glia and oligodendrocytes. The mixed culture was subsequently treated with a panel of drugs (Table 1) at 10 *µ*M for 24 h and frozen in a lysis buffer until library preparation. RNA was extracted and reverse transcribed into cDNA in each well of the plate, followed by pooling and preparation of mRNA libraries. After sequencing, mRNA reads were demultiplexed according to well barcodes, and the resulting gene expression profiles were processed by a standard differential expression method to derive drug-associated gene lists. **b**, A highlight of two compounds whose gene lists consistently yield improved performance over the randomly-selected lists of equal length. Shown is performance associated with predicting early-vs-late disease stages in several AMP-AD datasets. Each row corresponds to an evaluation of gene lists in a single dataset; MSBB evaluation is subdivided into four brain regions, specified as Brodmann Area. The vertical line denotes performance of the drug-associated list, while the background distribution shows performance of gene lists randomly selected from the same dataset. The drugs are annotated with their nominal targets.

### 3’ Digital Gene Expression profiles drug-induced perturbation of mRNA expression

Drug responses were generated using the RenCell VM human neural progenitor cell line. Upon growth factor withdrawal, RenCell VM cells differentiate into a mixed culture of neurons, glia and oligodendrocytes over a period of ∼7 days^37^. Differentiated RenCells were exposed to one of 80 drugs at two doses in triplicate, or to a vehicle-only (DMSO) control, for 24h and mRNA levels were then measured using the high-throughput, intermediate read density RNAseq method 3’ Digital Gene Expression (3’ DGE)^38^ (Fig. 2a); the advantage of DGE in this setting is that it provides high quality gene expression data at relatively low cost, allowing data to be collected from multiple repeats, doses and drugs. The drug panel was designed to include FDA-approved compounds, which could potentially be repurposed, compounds that have been tested in human clinical trials, and for which toxicity data are available, that could be further developed for AD and pre-clinical compounds designed to extend the range of targets and test therapeutic concepts not explorable with clinical grade compounds (Table 1). The 80 compounds were profiled across two separate 3’ DGE experiments and, to establish reproducibility, five of the 80 compounds were included in both experiments. The experiments are indexed separately (Table 1), and the compounds in common provide a measure of biological and technical variation. High concordance of the measurements between the two experiments was observed (Supplementary Fig. 2) suggesting that batch effects are not strong. Overall the drug-response data comprised 767 DGE gene expression profiles.

**Table 1:** Eighty compounds profiled in differentiated neuroprogenitor cell cultures. Each compound is annotated with its LINCS identifier, nominal target, approval status, toxicity, and strength of association with disease severity in ROSMAP and MSBB datasets (presented as HMP, harmonic mean p-value). Five of the compounds were profiled in two separate Digital Gene Expression (DGE) experiments.

| Plate | LINCSID | Drug | Target | Approval | IsToxic | HMP |
| --- | --- | --- | --- | --- | --- | --- |
| DGE1 | HMSL10024 | nvp-tae684 | ALK | experimental | 1 | 0.0015 |
| DGE1 | HMSL10045 | a443654 | AKT1 | experimental | 1 | 0.0023 |
| DGE1 | HMSL10177 | brivanib | VGFR1 | investigational | 0 | 0.0025 |
| DGE1 | HMSL10138 | ruxolitinib | JAK1 | approved | 0 | 0.0041 |
| DGE2 | HMSL10282 | vorinostat | HDAC1 | approved | 0 | 0.0047 |
| DGE2 | HMSL10127 | dovitinib | FLT3 | investigational | 0 | 0.0050 |
| DGE2 | HMSL10225 | regorafenib | c-Kit | approved | 0 | 0.0097 |
| DGE1 | HMSL10141 | fedratinib | JAK2 | investigational | 1 | 0.0115 |
| DGE1 | HMSL10226 | tofacitinib | JAK3 | approved | 0 | 0.0130 |
| DGE1 | HMSL10079 | torin1 | mTOR | experimental | 0 | 0.0170 |
| DGE2 | HMSL10159 | pelitinib | EGFR | investigational | 0 | 0.0186 |
| DGE2 | HMSL10099 | nilotinib | ABL1 | approved | 0 | 0.0249 |
| DGE2 | HMSL10051 | lapatinib | EGFR | approved | 0 | 0.0266 |
| DGE2 | HMSL10071 | palbociclib | CDK4 | approved | 0 | 0.0287 |
| DGE2 | HMSL10138 | ruxolitinib | JAK1 | approved | 0 | 0.0300 |
| DGE2 | HMSL10141 | fedratinib | JAK2 | investigational | 1 | 0.0304 |
| DGE2 | HMSL10411 | h89 | PKA | experimental | 0 | 0.0308 |
| DGE2 | HMSL10454 | staurosporine aglycone | NA | experimental | 0 | 0.0311 |
| DGE1 | HMSL10172 | gsk1059615 | PIK3CA | experimental | 1 | 0.0341 |
| DGE2 | HMSL10092 | jwe-035 | AURKA | experimental | 1 | 0.0373 |
| DGE1 | HMSL10194 | cabozantinib | VGFR2 | approved | 0 | 0.0399 |
| DGE2 | HMSL10213 | kin001-111 | SRC | experimental | 0 | 0.0440 |
| DGE1 | HMSL10192 | kw2449 | FLT3 | experimental | 0 | 0.0479 |
| DGE1 | HMSL10062 | gsk1070916 | AURKB | experimental | 0 | 0.0489 |
| DGE2 | HMSL10354 | baricitinib | JAK1/2 | approved | 0 | 0.0501 |
| DGE2 | HMSL10151 | nintedanib | VGFR1 | approved | 1 | 0.0594 |
| DGE2 | HMSL10226 | tofacitinib | JAK3 | approved | 0 | 0.0629 |
| DGE2 | HMSL10098 | gefitinib | EGFR | approved | 0 | 0.0650 |
| DGE1 | HMSL10055 | bx-912 | PDK1 | experimental | 0 | 0.0659 |
| DGE1 | HMSL10107 | mg-132 | PSB5 | experimental | 1 | 0.0665 |
| DGE1 | HMSL10582 | fccp | NA | experimental | 0 | 0.0730 |
| DGE2 | HMSL10139 | azd-1480 | JAK2 | investigational | 0 | 0.0740 |
| DGE1 | HMSL10311 | entinostat | HDAC1 | investigational | 0 | 0.0807 |
| DGE2 | HMSL10393 | at9283 | AURKA/B-JAK2 | investigational | 1 | 0.0837 |
| DGE2 | HMSL10318 | resveratrol | PGH1 | approved | 0 | 0.0864 |
| DGE2 | HMSL10020 | dasatinib | ABL1 | approved | 0 | 0.0867 |
| DGE2 | HMSL10310 | belinostat | HDAC1 | approved | 0 | 0.0899 |
| DGE2 | HMSL10228 | ikk16 | IKKA | experimental | 1 | 0.0964 |
| DGE1 | HMSL10241 | bortezomib | PSB5 | approved | 0 | 0.0978 |
| DGE1 | HMSL10096 | zm-447439 | AURKA | experimental | 0 | 0.0998 |
| DGE2 | HMSL10192 | kw2449 | FLT3 | experimental | 0 | 0.1148 |
| DGE2 | HMSL10069 | enzastaurin | KPCB | investigational | 0 | 0.1149 |
| DGE1 | HMSL10115 | ldn-193189 | ACVR1 | experimental | 1 | 0.1193 |
| DGE1 | HMSL10340 | hg-9-91-01 | SIK1 | experimental | 0 | 0.1199 |
| DGE2 | HMSL10394 | ceritinib | ALK | approved | 1 | 0.1218 |
| DGE2 | HMSL10091 | xmd16-144 | AURKA | experimental | 0 | 0.1282 |
| DGE2 | HMSL10150 | ponatinib | ABL1 | approved | 1 | 0.1283 |
| DGE1 | HMSL10328 | (+)-jq1 | BRD2 | experimental | 0 | 0.1361 |
| DGE2 | HMSL10232 | dactolisib | mTOR | investigational | 0 | 0.1448 |
| DGE2 | HMSL10050 | az-628 | BRAF | experimental | 0 | 0.1511 |
| DGE2 | HMSL10378 | ly2090314 | GSK3 | experimental | 0 | 0.1578 |
| DGE2 | HMSL10434 | bix 02565 | RSK | experimental | 0 | 0.1598 |
| DGE1 | HMSL10426 | barasertib | AURKB | investigational | 0 | 0.1704 |
| DGE2 | HMSL10440 | vx-702 | p38 MAPK | investigational | 0 | 0.1717 |
| DGE2 | HMSL10120 | canertinib | EGFR | investigational | 0 | 0.1864 |
| DGE2 | HMSL10080 | torin2 | mTOR | experimental | 1 | 0.2027 |
| DGE2 | HMSL10097 | erlotinib | EGFR | approved | 0 | 0.2216 |
| DGE1 | HMSL10049 | plx-4720 | BRAF | experimental | 0 | 0.2383 |
| DGE2 | HMSL10443 | axitinib | VEGFR | approved | 0 | 0.2397 |
| DGE1 | HMSL10441 | sb202190 | p38 MAPK | experimental | 0 | 0.2403 |
| DGE2 | HMSL10008 | sorafenib | BRAF | approved | 0 | 0.2480 |
| DGE2 | HMSL10027 | crizotinib | c-Met | approved | 1 | 0.2491 |
| DGE2 | HMSL10129 | ibrutinib | BTK | approved | 0 | 0.2549 |
| DGE2 | HMSL10364 | mrt67307 | TBK1 | experimental | 0 | 0.2571 |
| DGE2 | HMSL10018 | neratinib | ERBB2 | approved | 0 | 0.2669 |
| DGE1 | HMSL10165 | pf04217903 | c-Met | experimental | 0 | 0.2767 |
| DGE1 | HMSL10391 | alisertib | AURKA | investigational | 0 | 0.2900 |
| DGE2 | HMSL10117 | celastrol | PSB5 | experimental | 1 | 0.2940 |
| DGE2 | HMSL10133 | afatinib | ERBB2 | approved | 0 | 0.2987 |
| DGE1 | HMSL10470 | r 59949 | DGK | experimental | 0 | 0.2993 |
| DGE2 | HMSL10049 | plx-4720 | BRAF | experimental | 0 | 0.3213 |
| DGE1 | HMSL10125 | pha-665752 | c-Met | experimental | 0 | 0.3352 |
| DGE2 | HMSL10175 | sunitinib | VGFR1 | approved | 0 | 0.3557 |
| DGE2 | HMSL10284 | dabrafenib | BRAF | approved | 0 | 0.3693 |
| DGE2 | HMSL10189 | bosutinib | Src | approved | 1 | 0.4205 |
| DGE2 | HMSL10023 | imatinib | ABL1 | approved | 0 | 0.4262 |
| DGE1 | HMSL10114 | pazopanib | VGFR1 | approved | 0 | 0.4851 |
| DGE1 | HMSL10475 | r 59-022 | DGK | experimental | 0 | 0.4899 |
| DGE2 | HMSL10388 | xl019 | NA | investigational | 0 | 0.5485 |
| DGE2 | HMSL10198 | vandetanib | VGFR2 | approved | 0 | 0.6025 |
| DGE2 | HMSL10304 | rucaparib | PARP1 | approved | 0 | 0.6338 |
| DGE2 | HMSL10383 | pf-06463922 | ROS1 | experimental | 0 | 0.6371 |
| DGE1 | HMSL10586 | rotenone | NA | vet_approved | 0 | 0.6690 |
| DGE2 | HMSL10176 | y-27632 | ROCK1 | experimental | 0 | 0.7245 |
| DGE1 | HMSL10622 | metformin | NA | approved | 0 | 0.8090 |

For each drug, we defined the DGL to be the set of significantly perturbed genes, as identified through differential gene expression analysis^39^ comparing 3’ DGE profiles of drug-treated and control cells (see Methods). For most drugs, we identified between 10 and 300 significantly perturbed genes. DRIAD evaluated each DGL by constructing a predictor of pathological stage based on mRNA expression of these genes in AMP-AD datasets. The accuracy of the predictor assessed over multiple brain regions measures the association between a drug’s “mechanism” (encapsulated in the DGL) and AD severity. We focused specifically on the binary classification task of distinguishing early vs. late disease stages (A-vs-C), because it contrasts groups of maximally distinct samples and yields higher signal to noise ratio than attempting to predict all three disease stages. This in turn leads to higher predictive power of random gene lists and a higher performance bar that DGLs will need to surpass to be considered significant.

### Systematic assessment of drug signatures derived from 3’ DGE leads to a ranked list of repurposing candidates

Each DGL is evaluated against randomly-selected gene lists of the same length (Fig. 1a). The empirical p value, computed as the fraction of random lists that outperform a DGL, constitutes a natural starting point for comparing predictor performance across drugs, because it is effectively normalized by the number of genes in the DGL. An example of drugs whose DGL consistently outperform random lists include the pre-clinical compound TAE684 and the approved drug ruxolitinib, whose primary targets are ALK and JAK1/2 kinases respectively (Fig. 2b). Next, we combined the empirical p values from various datasets and brain regions to create a harmonic mean p-value (HMP) for each drug. The HMP facilitates the detection of significant hypothesis groups in a multiple hypothesis setting, while being less restrictive and providing greater statistical power than similar multiple hypothesis testing procedures^40^. Using HMP as a prioritization scheme, we identified the top 15 FDA-approved and top 15 pre-clinical drugs (Fig. 3) in the full ranking of all 80 drugs that were profiled in neuronal cultures (Table 1).

**Fig. 3:**
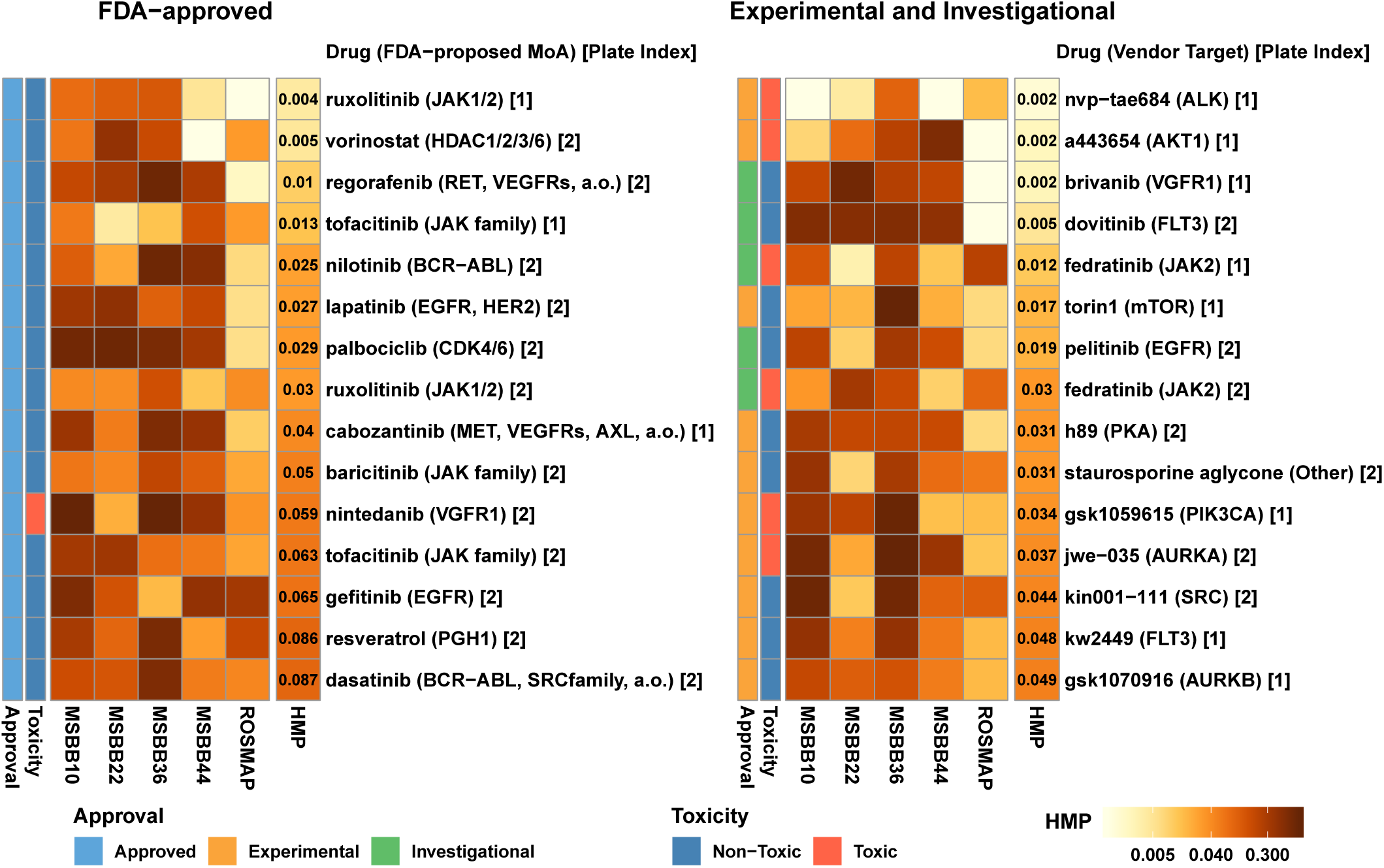
Top 15 FDA-approved (left) and experimental/investigational (right) drugs, sorted by harmonic mean p-value. Each heatmap shows p-values associated with a drugs predictive performance across two AMP-AD datasets, ROSMAP and MSBB. The MSBB analysis is further subdivided by the brain region, specified as Brodmann Area. The last column in each heatmap shows the harmonic mean p-value (HMP). The rows are annotated with the name of the drug/compound, its nominal target, and the index of the corresponding DGE experiment. Additional annotations include information about each compounds approval status (approved / investigational / experimental) and whether compounds were found to be toxic in neuronal cell cultures.

Some of top-performing drugs were associated with drugs that are known to be cytotoxic (e.g. TAE684^41^) and were discovered as antineoplastics. To assess the magnitude of this cytotoxicity, we quantified viable differentiated RenCells following 48 h of exposure to a high drug dose (10 uM) using fixed cell microscopy (see Methods and Supplementary Fig. 3); drugs that significantly reduced viable cell number were annotated cytotoxic (Fig. 3). A general consensus was observed between viable cell number and the total RNA yield (Wilcoxon Rank Sum test, p < 4 × 10^−9^), suggesting that compound toxicity can also be inferred directly from a reduction in mRNA abundance in post-perturbational gene expression profiles (Supplementary Fig. 3). Since DGLs of cytotoxic drugs consistently outperform random lists across multiple AMP-AD dataset (Fig. 2b, 3), the mechanisms of cell death induced by these drugs may share some similarity with mechanisms of cell death in AD. Screening for drugs that rescue death induced by TAE684 (or other cytotoxic compounds) might therefore identify compounds with the ability to prevent cell death of human neural cells but such a screen is outside of the scope of the current work.

### Elucidating target affinity spectrum properties associated with the observed drug ranking

Do high-scoring drugs have features in common? With respect to primary targets, we observed that many drugs, including ruxolitinib, inhibit one or more of members of Janus Kinase family, which comprises JAK1, JAK2, JAK3 and TYK2 (Fig. 3). However, these compounds are known to have additional off-targets that might also contribute to activity^19,42^. To investigate the potential role of these secondary targets we used the Target Affinity Spectrum (TAS), a vector computed from experimental data that quantifies the potency of a drug against a range of targets^19^. TAS vectors were constructed by aggregating information about targets and non-targets from published dose-response measurements, experimental profiling data, and manual annotations in the literature. Confirmed drug-target bindings were given a TAS score of 1,2, or 3 (with lower values indicating higher binding affinity), while confirmed non-binders were annotated with a TAS score of 10 (Fig. 4a). The full set of TAS scores, which includes the 80 compounds considered in this study, is publicly available at http://smallmoleculesuite.org.

**Fig. 4:**
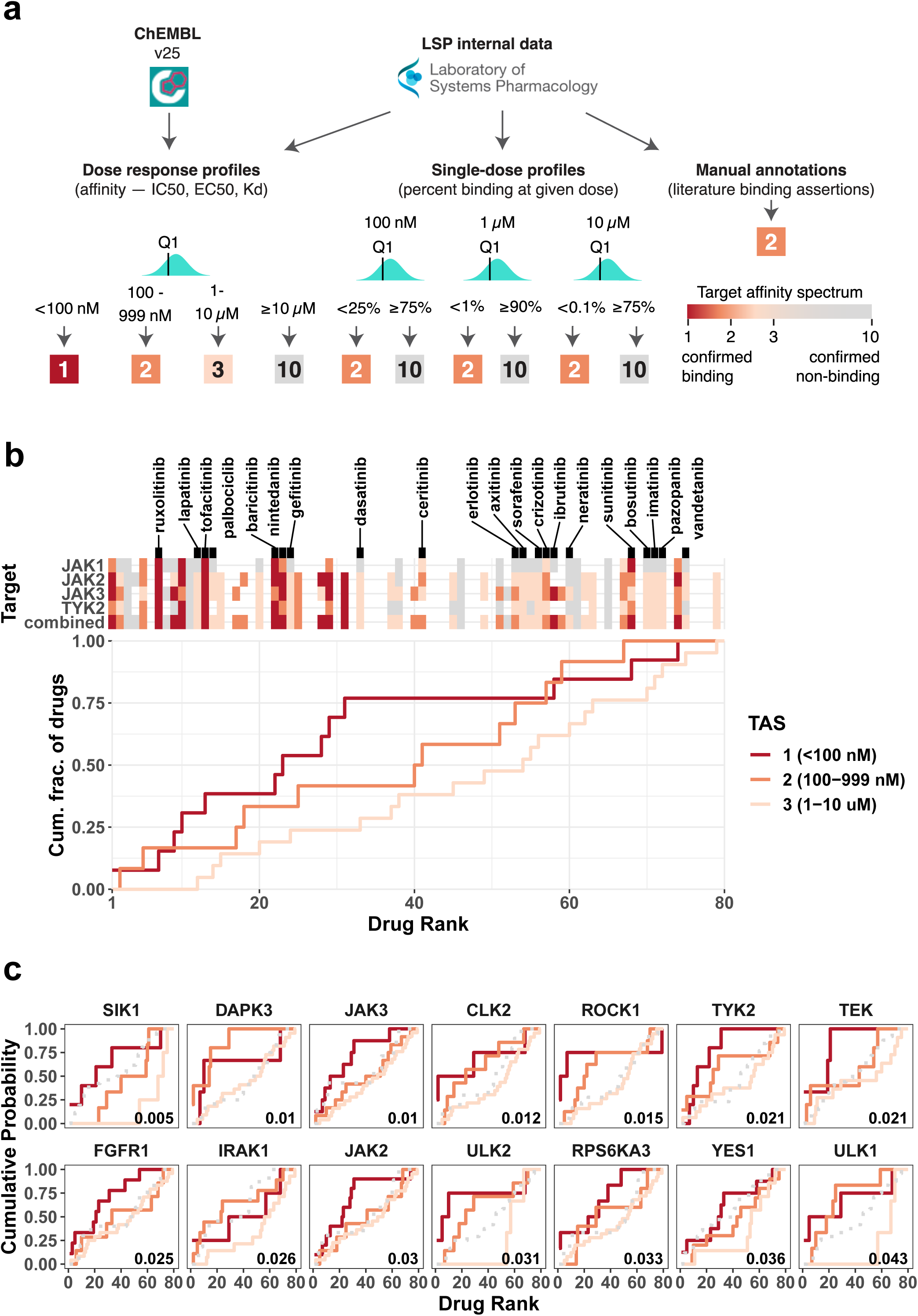
**a**, Overview of Target Affinity Spectrum (TAS) score computation from raw drug binding data. Three types of drug binding data were sourced from ChEMBL and from the internal Laboratory Systems of Pharmacology dataset that have not yet been incorporated into ChEMBL. Empirically derived thresholds for the different data types where used to assign TAS scores to each drug-target pair. Multiple measurements for the same drug-target combination were aggregated along the first quartile to define the final TAS value. **b**, Binding affinity of compounds in the ranked list to every member of the Janus Kinase family. The compounds are sorted in increasing order by the harmonic mean p-value (as defined in Figure 3) along the x-axis. The top heatmap shows the binding affinity of each compound to the selected targets, explicitly naming the FDA-approved drugs. Colored and gray tiles denote confirmed binders and non-binders, respectively; missing entries correspond to unknown affinity values. The combined affinity is defined as the strongest binding (lowest TAS score) among all four JAK targets. The bottom plot shows the breakdown of the combined affinity values by TAS-specific empirical cumulative distribution functions (ecdfs). Each line shows ecdfs for all drugs that bind the corresponding target with a TAS score of 1 (dark orange), 2 (orange) or 3 (light orange). **c**, Top targets whose binding affinity correlates most strongly with the compound ranking. The ecdfs of confirmed non-binders (TAS = 10) are shown as gray dashed lines for reference. Area under ecdf can be interpreted as a summary statistic that captures the position of drugs binding to that target with the corresponding affinity in the ranked list. Correlation between the drug ranking and TAS values was computed using Kendalls Tau test, with the associated p value displayed in the bottom right corner of each plot.

We evaluated whether the strength of a binding interaction (i.e., binding affinity) between a compound and its target or a set of biologically related targets contributes to the ranking of a compound by DRIAD. Significant positive correlations between the drug ranking and the binding affinity suggest that the target or a class of targets are likely to be pertinent to one or more of the disease mechanisms. To illustrate this effect, we constructed empirical cumulative distribution functions (ECDFs) for the binding of drugs to members of the JAK family. The list of 80 drugs was traversed in the order of increasing harmonic p-value, while keeping track of the cumulative count of drugs with a particular TAS value (Fig. 4b). This results in three different ECDFs, representing all drugs that bind to the corresponding target with a TAS score of 1, 2, or 3. Area under individual ECDF curves (AUC) can be interpreted as a summary statistic capturing the position of JAK binders with a particular affinity in the ranking; larger values of the AUC correspond to a higher saturation of the corresponding drug set near the top of the ranking.

We observed that compounds having higher affinity for members of the JAK family (i.e., lower TAS values) tend to appear earlier in the ranking (Fig. 4b), with a significant correlation (p = 0.001; Kendall’s Tau test) between binding affinity and the drug ranking as defined by DRIAD. This suggests that downstream transcriptional changes induced by JAK inhibitors correlate with Braak AD stage severity *in an affinity-dependent manner*. Direct inspection of the experimental 3’ DGE data confirms that binding affinity directly correlates with the level of expression in several interferon-stimulated genes, with drugs that have a higher affinity to JAK family members resulting in stronger inhibition of interferon gene expression (Supplementary Fig. 4). These observations build on previous studies^43^ suggesting that inhibition of the JAK-STAT and interferon signaling pathways might be beneficial in the context of AD.

We repeated the binding affinity correlation analysis for all targets that had confirmed binding interactions with at least three drugs used in this study (Fig. 4c). The results showed that most of the affinity-dependent effects were contributed by “off-targets”, i.e., established targets for a drug that are different from the target for which the drugs is marketed (the ‘nominal target’). For example, we observed strong associations between drug ranking and binding affinity for Unc-51-Like Kinases 1 and 2 (ULK1, ULK2) and their downstream substrate Death-Associated Protein Kinase 3 (DAPK3), all of which are associated with autophagy^44–46^. Autophagy plays an important role in cellular homeostasis and its dysregulation is emerging as a contributing factor to neurodegenerative diseases, including AD^47,48^. Previous studies suggest that inhibition of autophagy may impair clearance of neurotoxic aggregates, and future effective therapy might require that the level of autophagy is maintained, or even increased, as part of upstream perturbations^49,50^. The Salt Inducible Kinase 1 (SIK1) also appears to have a strong association with the position of compounds in the ranking. This association is driven primarily by TAE684, fedratinib, and GSK1070916, all of which are strong binders of SIK1 (TAS = 1) and appear near the top of ranking established by DRIAD. However, none of these drugs are FDA approved or have been studied in the context of SIK1 inhibition.

### Polypharmacology analysis reveals additional mechanisms that may correlate with AD severity

We also considered downstream effects of concerted off target binding (polypharmacology) in which genes with a closer association to disease severity are altered more significantly by coordinated activity on two or more targets relative to drugs that selectively bind to only one of the off-targets. To evaluate the impact of polypharmacology on drug-disease associations, we divided the 80 compounds in our dataset into three categories: those with confirmed binding to Target A and Target B and those that bind either Target A or Target B alone (see Methods). The three categories were then compared to determine whether compounds binding both targets appear significantly closer to the top of the ranking (Fig. 5a) as defined by the harmonic mean p-value computed by DRIAD (Table 1). We systematically evaluated all pairs of targets that had a least six compounds with confirmed binding interactions associated with TAS values of 1, 2, or 3, and then identified the top pairs in the ranking (Fig. 5b). A pair was deemed to be a positive interaction if compounds binding both targets (A and B) appeared significantly closer to the top of the ranking than compounds binding only one of the targets (A or B, but not both); the pair was deemed to be a negative interaction if the opposite held true.

**Fig. 5:**
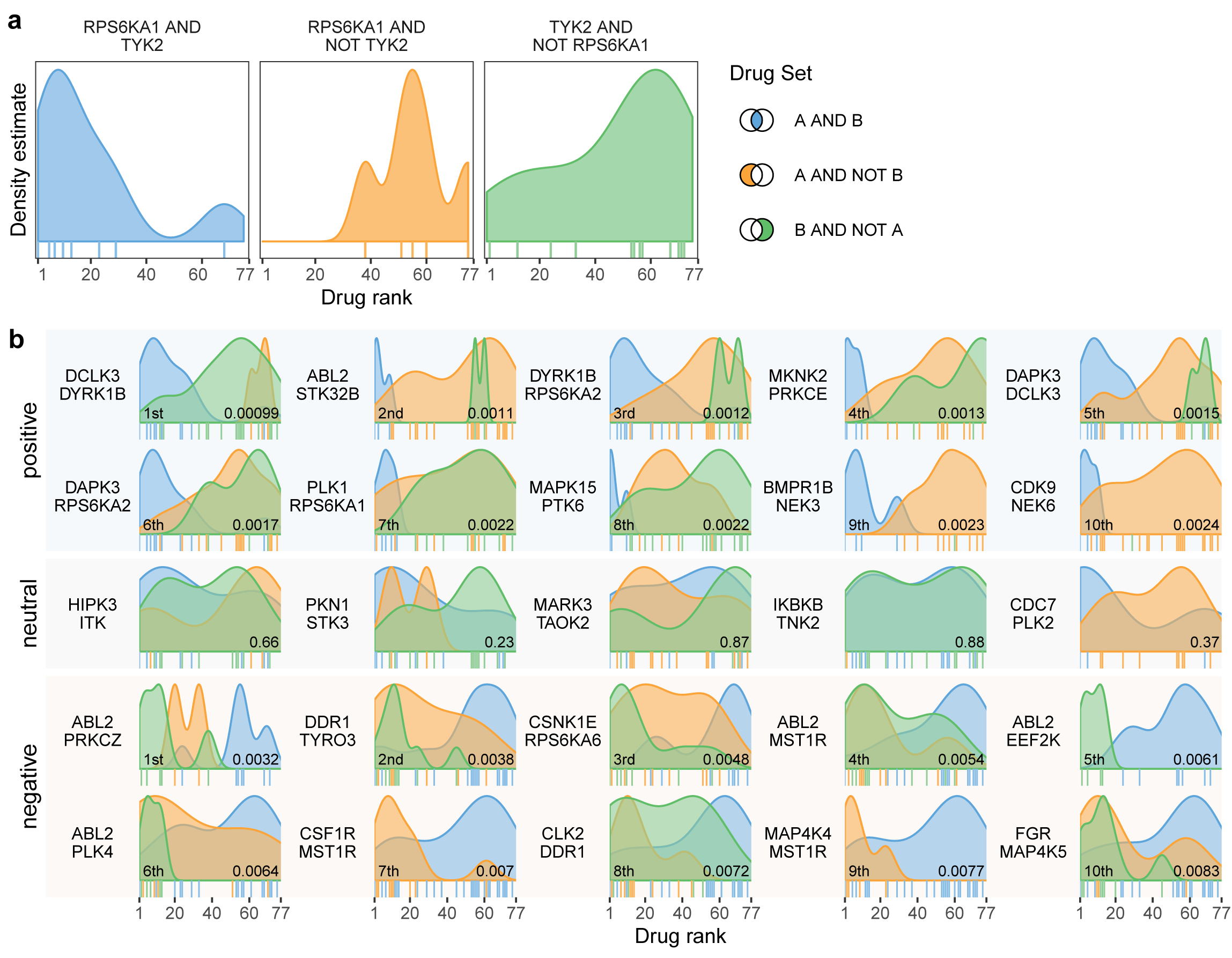
**a**, An example polypharmacology test with a focus on RPS6KA1 and TYK2. The drugs are ranked by the harmonic mean p-value (as in Figures 3 and 4), and the distributions of drugs bindings to both RPS6KA1 and TYK2 (left), those binding to RPS6KA1 but not TYK2 (middle) and, conversely, TYK2 but not RPS6KA1 (right) are shown along this ranking. Individual drugs that bind those targets are annotated by vertical tick marks directly below the corresponding distribution. **b**, Top 10 synergistic and top 10 antagonistic relationships between pairs of targets. The distributions in each plot are compared using Wilcoxon Rank Sum test, with the resulting p value presented in the bottom right corner. If compounds that bind both targets appear significantly closer to the top of the ranked list (left side of the x axis), we define the target pair to be synergistic. Conversely, a pair of targets with an explicit non-binding interaction observed among the top-ranking compounds is defined to be antagonistic. A set of five neutral target pairs (i.e., no significant synergistic or antagonistic effect) is included for reference.

To determine whether one or more targets consistently participate in positive or negative interactions with other targets we aggregated individual p values from the evaluation of pairs that contained the target of interest, using the Brown’s method and Jaccard similarity as the metric of independence between individual tests (see Methods). The list of targets was subsequently sorted by the aggregated p value (Table 2 and Supplementary Table 1). We found that several gene families emerged as candidate for top scoring compounds in which poly-pharmacology was predicted to be essential. For example, the top-scoring compounds TAE684, dovitinib, ruxolitinib and fedratinib are observed to bind RPS6KA1 (Fig. 5a, Table 2), a component of a microglial signature^51^ with a potential role in AD as identified by previous epigenetic studies^52^, and RPS6KA2, a gene involved in Neurotrophin signaling^53^ with previous reports of association with Parkinson’s Disease in GWAS studies^54^. Similarly, NIMA-related Kinases NEK3, NEK6 and NEK9 consistently appear in positive interaction with other drug targets among top-scoring compounds (Table 2). All three genes have known relationships to microtubule function and Tau phosphorylation. NEK3 has been reported to influence neuronal morphogenesis through microtubule acetylation, and its phospho-defective mutant is hypothesized to play a role in axonal degeneration^55^. NEK6 phosphorylates p70-S6^56^, a key player in hyperphosphorylation of Tau^57,58^ that leads to microtubule disruption and deposition of Tau tangles. It was found to be relevant to the progression of AD and proposed as an early diagnostic biomarker^59,60^ NEK9 was found to be differentially expressed in a tauopathy mouse model^61^. Taken together, these results suggest that the NEK family may be an important set of co-targets, and a successful future therapeutic might require polypharmacology with respect to these kinases.

**Table 2:** Top 10 targets that consistently appear in the top positive or negative interactions. The table lists targets, the number of pairs they appear in, whether those pairs are primarily positive or negative interactions, and the overall p-value computed by aggregating p-values from individual Wilcoxon Rank Sum tests (Fig. 5b) using the Brown’s method.

| symbol | direction | n | p | padj |
| --- | --- | --- | --- | --- |
| NEK6 | positive | 257 | 8.25E-129 | 1.32E-06 |
| NEK3 | positive | 237 | 5.45E-71 | 2.52E-04 |
| RPS6KA2 | positive | 246 | 7.62E-68 | 4.80E-04 |
| LATS2 | positive | 280 | 3.82E-60 | 9.53E-04 |
| ABL2 | negative | 204 | 2.88E-51 | 9.94E-04 |
| DCLK3 | positive | 250 | 1.23E-56 | 0.00124436 |
| MARK1 | positive | 271 | 4.12E-52 | 0.00191891 |
| STK17B | positive | 264 | 1.22E-49 | 0.00221427 |
| NEK9 | positive | 295 | 5.14E-50 | 0.00266415 |
| STK17A | positive | 202 | 8.57E-37 | 0.0032299 |

## Discussion

In this paper we described the development of DRIAD, a machine learning framework for evaluating potential relationships between a disease and any biological process than can be described by a list of genes. We used DRIAD to look for associations between the pathological stage of AD and genes that are differentially expressed in neurons exposed to potential therapeutic drugs. DRIAD is distinct from the traditional approaches in which a model is constructed over the entire gene space and subsequently interrogated for feature importance scores and the enrichment of predefined gene sets, which then serve as a candidate list for further functional studies^62^. The traditional approach is well-suited for predictors that exhibit high accuracy, because they establish a strong association between input features and the predicted phenotype. As predictor accuracy decreases however, it becomes difficult to distinguish whether a high enrichment score is a true association between the corresponding mechanism and disease, or an artifact in a model that doesn’t accurately predict the phenotype of interest (here, disease stage as defined by Braak score). DRIAD effectively decouples gene set enrichment and predictor performance by pre-filtering the transcriptomic space to genes associated with drugs prior to model training and predictor evaluation. Pre-filtering to a limited set of features also addresses the issue of overfitting that often arises when constructing computational models from disease databases where the number of cases is much less than the number of ‘omic features. Thus, DRIAD enables a direct, unbiased quantification of the association between drugs and AD.

The eighty compounds that we profiled in vitro were predominantly kinase inhibitors with anti-cancer activity since this is the largest class of targeted drugs currently available, both as approved and pre-clinical compounds^19^, with extensive target information. In addition, there is an inverse relationship between incidence of cancer and incidence of Alzheimer’s disease^63,64^. Of the 80 compounds tested, 35 were FDA-approved (Table 1) and can potentially be used for repurposing. The remaining set consisted of 35 pre-clinical and 15 investigational compounds, which allowed us to explore a wider spectrum of mechanisms. Targets of pre-clinical and investigational compounds that were scored highly by DRIAD could potentially be used for selection of FDA-approved compounds in future screens or for the development of NMEs.

We ranked all compounds by how well their mechanism of actions (as represented by a list of gene names) were able to predict disease severity based on gene expression in AMP-AD datasets. We found several drugs whose primary targets are JAK kinases to be among the top performers. We also explored connections between drug and their primary and secondary targets. This revealed that top-rankings drugs modulate pathways related to interferon signaling, autophagy and microtubule formation and function. Kinases from JAK, ULK and NEK families were found to be among the most consistent targets of top-scoring drugs. Future investigation will include experimental validation of these targets in cell-based and animal models using CRISPRa and/or CRISPRi, or other gene editing techniques, to evaluate whether a drug “hit” from DRIAD has an impact on AD-associated pathophysiology.

DRIAD has the potential to identify drugs that both mimic (or accelerate) disease and those that inhibit it. From the perspective of actual drug repurposing only the later compounds are useful. In other studies, gene expression changes have demonstrated increased interferon signaling in AD^65^ and in ALS^66^ brains. Recently, we have found that cytoplasmic dsRNA, a known activator of Type I interferon, is present in ALS brains with TDP-43 pathology. Similarly, cytoplasmic dsRNA, which has been linked to increased Type I interferon signaling, was found to accumulate in glia in AD brains^67^. Activation of interferon signaling in this context promotes neuronal cell death. Thus, it seems probable that the inhibitors of JAK-STAT signaling identified in this study will potentially be useful in blocking neuroinflammation and cell death in the context of AD. Further studies to investigate the role of JAK-STAT signaling in aging and in AD brains are therefore warranted.

DRIAD allows for unbiased assessment of biological processes or drug candidates even when disease mechanisms are not explicitly known. This is valuable from a neuropathological perspective because it is increasingly clear that in addition to the classical AD hallmarks of amyloid plaques, neurofibrillary tangles, and neuronal loss, most patients with a clinical diagnosis of AD dementia have distinct patterns of co-occurring pathologies including TDP43 inclusions, Lewy bodies, vascular changes, astrocyte and microglial activation, and probably other unrecognized alterations^22,68–71^. By working directly with the mRNA expression data from postmortem brain specimens and *a priori* knowledge of which genes encapsulate a proposed mechanism or co-expression module, or which genes are perturbed by a set of drugs, DRIAD can score mechanisms, co-expression modules or drugs without explicit knowledge of co-existing pathologies, e.g. presence of Lewy Bodies or TDP-43 inclusions, in individual patients. Thus, DRIAD is capable of evaluating diverse hypotheses, including those associated with repurposing, without a high level of prior knowledge.

In the datasets used for this study, gene expression profiles from autopsied brains are associated with Braak pathologic stage, making it possible to compare patients with no or mild AD symptoms at the time of death to patients who were demented. However, this anchoring on Braak staging also includes some cases where symptoms did not correlate with pathological stage. A follow-on computational approach to deconvolve the correlative signals observed between top-performing drug signatures and AD expression profiles would help inform subsequent mechanistic studies. One direction for follow up is to rerun the DRIAD pipeline on patient subgroups as defined by more detailed clinical or pathologic phenotypes, motivated by the notion of personalized treatment: different molecular pathways of AD will likely require different interventions to rescue neuronal death.

This study has identified associations of gene perturbations by FDA-approved drugs and investigational compounds in human neural cells with gene perturbations in AD brain regions. Our results require validation in relevant model systems or through emulated clinical trials in electronic health records^72^. Parallel mechanistic studies to discover relevant CNS-specific pharmacodynamic biomarker(s) for a drug action, in the cerebrospinal fluid, for instance, would enable formal evaluation of each drug’s efficacy and safety in randomized clinical trials.

## Methods

### High-Throughput profiling using 3’ Digital Gene Expression

A multiwell cell dispenser (catalog# 5840300, Thermo Scientific, Waltham, MA) with standard tubing (catalog# 24072670, Thermo Scientific, Waltham, MA) was used to plate 2500 neural stem cells (ReNcell VM, catalog# SCC008, Millipore, Billerica, MA) into each well of a 384-well cell culture plate (Perkin Elmer, Waltham, MA). Neural stem cells where differentiated into mature neural cells for one week and then treated with compounds (Table 1) or DMSO using a D300 Digital Dispenser (Hewlett-Packard, Palo Alto, CA). D300 software was used to randomize dispensation of compounds. After 24 hours, the cells were washed once with PBS using an EL405x plate washer (BioTek, Winooski, VT) leaving 5-10 ul of PBS behind in each well. 10 ul of 1X TCL lysis buffer (catalog# 1070498, Qiagen, Hilden, Germany) with 1% (v/v) ß-mercaptoethanol was added per well, and the plates were stored at -80°C until the RNA extraction was performed.

For RNA extraction, the cell lysate plate was thawed and centrifuged for 1 min at 1000 rpm. Using a BRAVO (Agilent, Santa Clara, CA) liquid handler, the lysate was mixed thoroughly before transferring 10 ul to a 384 well PCR plate. 28 ul of home-made Serapure SPRI beads (GE Healthcare Life Sciences, Marlborough, MA) were added directly to the lysate, mixed and incubated for 5 min. The plate was transferred to a magnetic rack and incubated for 5 min prior to removing the liquid to aggregate the beads. The beads were washed with 80% ethanol twice, allowed to dry for 1 min, 20 ul of nuclease free water was added per well, the plate was removed from the magnetic rack and the beads were thoroughly resuspended. Following a 5 min incubation, the plate was returned to the magnetic rack and incubated an additional 5 min before transferring the supernatant to a fresh PCR plate. 5 ul of the RNA was transferred to a separate plate containing RT master mix and 3’ and 5’ adapters for reverse transcription and template switching (Soumillon et al., 2014), and incubated for 90 min at 42°C. The cDNA was pooled and purified with a QIAquick PCR purification kit according to the manufacturer’s directions with the final elution in 21 ul of nuclease free water. This was followed by exonuclease I treatment for 30 min at 37°C that was stopped with a 20 min incubation at 80°C. The cDNA was then amplified using the Advantage 2 PCR Enzyme System (Takara, Fremont, CA) for 6 cycles, and purified using AMPure XP magnetic beads (Beckman Coulter Genomics, Chaska, MN). Library preparation was performed using a Nextera XT DNA kit (Illumina, San Diego, CA) on 5 reactions per sample following the manufacturer’s instructions, amplified 12 cycles, and purified with AMPure XP magnetic beads (Beckman Coulter Genomics, Chaska, MN). The sample was then quantified by qPCR and sequenced on a single Illumina NextSeq run with 75bp paired end reads at the Harvard University Bauer Core Facility.

Raw RNA reads were aligned against a reference genome and quantified using the bcbio-nextgen single cell/DGE RNA-seq analysis pipeline (https://bcbio-nextgen.readthedocs.io/). The pipeline consists of the following steps: 1) well barcodes and unique molecular identifiers (UMIs) were extracted from every RNA read; 2) all reads not within the edit distance of a single nucleotide from a predefined well barcode were discarded; 3) each extant read was quasialigned to the human transcriptome (version GRCh38) using RapMap^73^; 4) reads per well were counted according to UMIs^74^, discarding reads with duplicate UMIs, weighting multi-mapped reads by the number of transcripts they aligned to and collapsing transcript counts to gene level by summing across all transcripts of a gene.

Differential gene expression analysis was performed with the R package edgeR 3.18.1. Compound-associated gene lists were composed from genes with a significant (FDR < 0.05) post-perturbation change in expression level compared to DMSO controls. List length was capped at 300 genes to ensure that <2% of the transcriptional space is captured by any one compound-associated list, thus maintaining a wide diversity among randomly-selected lists of matching lengths.

### Prediction of disease stage

Gene expression profiles of postmortem brain specimens along with the corresponding clinical annotations were downloaded from the Synapse portal at www.synapse.org/AMPAD. The entire transcriptional feature space was filtered down to ∼20k protein-coding genes to ensure fairness of comparison between transcriptional changes induced by the profiled compounds (which were all kinase inhibitors in this study) and the randomly-sampled background lists. Every specimen was assigned a label of disease severity based on the following mapping to the Braak annotations^18^: A - early (Braak 0-2), B - intermediate (Braak 3-4), and C - late (Braak 5-6). The labels were used to establish three binary classification tasks, contrasting A vs. B, B vs. C, and A vs. C, with the expression of a prespecified list of genes playing the role of the input features.

Data for any given binary classification task were used to train a logistic regression classifier, which models log-odds ratio between the two classes as a linear combination of the input features. Logistic regression was chosen due to its popular use with gene expression data and because it guarantees that every preselected input feature participates in the assignment of predicted labels to new data points. (This is in contrast to methods such as random forests, where one or more features may not get selected to be included in at least one decision tree.) To address overfitting, a ridge regularization term that penalizes the L2-norm of feature weights was included in the model. No LASSO regularization was used, as it induces sparsity and excludes features that were specifically preselected to be included in the model.

Note that the proposed machine learning methodology is fairly general and, ultimately, the choice of a classification method does not matter, as long as the same predictor type is applied to random lists and the feature lists of interest. The desired properties are 1) that every feature in the list of interest is included in the model, and 2) significantly better than random predictor performance is obtainable with a relatively small number of features, thus allowing for an effective comparison of feature lists.

Model performance was evaluated through leave-pair-out cross-validation. For a given binary classification task, each example in the dataset was associated with the example from the opposite class that was the closest match in age. If there were multiple candidates for the age match, the pairing was selected uniformly at random. The resulting set of age-matched pairs was evaluated in a standard cross-validation setting, by asking whether the later-stage example in each withheld pair was correctly assigned a higher score by the corresponding predictor. The fraction of correctly-ranked pairs constitutes an estimate of the area under the ROC curve^75^.

### Assessing gene list significance

For a given gene list of interest, 1,000 random gene lists of matching cardinalities were sampled from a uniform distribution over the protein-coding space. Additional analysis did not reveal any significant association of predictor performance to the pairwise correlations among selected genes, nor to the proximity of selected genes on a gene-gene interaction network. Based on these observations, we saw no reason to bias random gene selection towards more (or less) internal connectivity.

Gene lists produced by the random sampling constitute a background for comparison with a particular gene list of interest. After evaluating all lists through cross-validation, an empirical p value was computed as the fraction of background lists that yield higher predictor performance than the gene list of interest. P values calculated for the same gene list across multiple datasets were combined to produce the Harmonic Mean P (HMP) value^40^. Gene lists associated with post-perturbational transcriptional changes were sorted by HMP to produce the final ranking of the corresponding compounds.

### Assessing toxicity

A multiwell cell dispenser (catalog# 5840300, Thermo Scientific,Waltham, MA) with standard tubing (catalog# 24072670, ThermoFisher Scientific,Waltham, MA) was used to plate 10,000 cells per well of a 96-well cell culture plate (catalog# 3603, Corning, Corning NY). Cells were treated with compounds (Table 1) or DMSO using a D300 Digital Dispenser (Hewlett-Packard, Palo Alto, CA). D300 software was used to randomize dispensation of compounds. After 48 hours, 60µl of a solution containing 10% Optiprep (catalog# D1556, Millipore Sigma, St. Louis MO) diluted with PBS (catalog# 21-040-CV, Corning, Corning NY), and a 1:5000 dilution of Hoechst 33342 (catalog# H3570, ThermoFisher Scientific, Waltham, MA) was gently added to the side of each well using a multichannel pipette (catalog# 1060-0850, VistaLab Technologies, Brewster, NY) on the lowest speed setting of 1. After 30 minutes, 80µl of a solution containing 3.7% formaldehyde (catalog# 15711, Electron Microsopy Sciences, Hatfield, PA), 20% Optiprep in PBS was added. After a 30-minute incubation a multichannel pipette (catalog# 1060-0850, VistaLab Technologies, Brewster, NY) was used to remove all but 15µl from each well. 100µl of 1xPBS was added to each well and the plate was covered with a foil seal (catalog# MSF-1001, Bio-Rad, Hercules, CA). Images were taken on an InCell 6000(GE Healthcare Bio-Sciences, Pittsburgh, PA). Columbus Image Data Storage and Analysis System (Perkin Elmer, Waltham MA) was used to count the number of Hoechst stained nuclei as a readout of cell number.

### Target affinity spectrum, top targets and polypharmacology analysis

#### Systematic classification of compound-target affinities using Target Affinity Spectrum (TAS)

TAS scores were calculated as described in the original paper^19^. Briefly, drug affinity data from ChEMBL v25^76^ and in-house data comprising drug affinity curves, single-dose binding data from the DiscoverX platform and manual binding assertions curated from literature were compiled into a single consistent measure of binding affinity. Multiple measurements for the same drug-target combination were aggregated by calculating the first quartile. For each drug-target pair, we only considered the highest quality source of data. If full dose-response affinity measurements were available, they took precedence over single-dose binding measurements, which took precedence over binding assertions mined from the literature.

Dose-response affinity data were converted to TAS scores based on empirically derived concentration cutoffs (<100nM: TAS=1; 100-999nM: TAS=2; 1-10µM: TAS=3 and >10µM: TAS=10). For single-dose drug binding data, we used concentration-specific thresholds derived from the empirical correlation between dissociation constant and percent inhibition (100nM <25%: TAS=2, ≥75%: TAS=10; 1µM <1%: TAS=2, ≥90%: TAS=10; 10µM <0.1%: TAS=2, ≥75%: TAS=10). Drug-target pairs were assigned TAS=2 or TAS=10 if they were mentioned in confirming (e.g. “drug X was equipotent for Y”) or negative (e.g. ‘‘drug X was found to not inhibit Y’’) statements in the literature.

All TAS values used in this study are publicly available through https://smallmoleculesuite.org.

#### Identification of important target genes using TAS profiles

Drugs were ranked based on their HMP scores, as computed above. If a drug was profiled in more than one 3’-DGE experiment, the corresponding HMP scores were averaged with a geometric mean. For a given target of interest, empirical cumulative distribution functions (ECDFs) were computed for each TAS value separately, using the HMP-based ranking as input. Area under individual ECDFs provides a summary statistic for the overall placement in the ranking of drugs with the corresponding TAS value.

The importance of a particular drug target was assessed through Kendall’s Tau test, which compares whether pairs of drugs are ordered the same way in two different rankings. In our case, a pair of drugs is considered to be concordant if the drug with the higher binding affinity (lower TAS value) appears closer to the front of the ranking, and discordant otherwise. The Kendall’s Tau coefficient is then defined as the fraction of concordant pairs among those that can be ordered (i.e., pairs of drugs with non-identical TAS values). The associated p-value of the Tau coefficient is approximated through standard permutation testing. Targets with fewer than three confirmed binders (TAS = 1,2,3) were not evaluated.

#### Polypharmacology analysis of target gene combinations

We first compiled a list of pairwise gene combinations for which we had TAS scores for both targets from at least six compounds. For each combination of targets, we split the compounds into three categories based on their TAS scores: compounds that bind both targets (category “A AND B”) and compounds that bind one of the targets but not the other (categories “A AND NOT B” and “B AND NOT A”). A union of the latter two was defined to be an “A XOR B” set. Wilcoxon rank sum tests were performed separately for “A AND NOT B”, “B AND NOT A”, and “A XOR B”. In all cases, the comparison was made relative to “A AND B”.

For each target pair, p-values from individual Wilcoxon rank sum tests were averaged using the harmonic mean^40^. The resulting p values were further aggregated using the Brown’s method (an extension of the Fisher’s method) with the test dependence metric being defined as the Jaccard similarity of the corresponding compound sets. For example, DCLK3 participated in 250 pair evaluations. Two of those evaluations included positive interactions “DCLK3 AND DYRK1B” and “DCLK3 AND DAPK3” pairings, with the corresponding harmonic mean p values 0.00099 and 0.0015, respectively (Figure 5B). There are ten compounds binding DCLK3 and DYRK1B, and the same ten compounds also bind to DCLK3 and DAPK3, yielding a Jaccard similarity of 1.0 for the two pairings. The two p values are thus considered to be coming from entirely non-independent tests by the Brown’s method, which aggregates all 250 p values into a single metric of importance for DCLK3 (Figure 5C).

### Data and code availability

Raw post-perturbational gene expression data for the 80 compounds profiled in this study, the associated gene lists, drug toxicity data and all relevant metadata have been uploaded to Synapse at http://synapse.org/DRIAD. The machine learning framework for evaluating the capacity of gene lists to predict disease severity is publicly available on GitHub at https://github.com/labsyspharm/DRIAD. Upon accepting the appropriate AMP-AD data usage agreements, users can evaluate their own gene lists on the different datasets, brain regions and binary classification tasks. Scripts to fully reproduce the tables and figures presented in this manuscript are also provided on GitHub at https://github.com/labsyspharm/DRIADrc. The reproducibility was made possible in part by the R packages grImport2^77^ and gridSVG^78^.

## Acknowledgments

The results published here are in part based on data obtained from the AMP-AD Knowledge Portal (doi:10.7303/syn2580853). These data were generated from postmortem brain tissue collected through the Mount Sinai VA Medical Center Brain Bank, led by Dr. Eric Schadt from Mount Sinai School of Medicine, by the Rush Alzheimer’s Disease Center, Rush University Medical Center, Chicago, and by the following sources: The Mayo Clinic Alzheimer’s Disease Genetic Studies, led by Dr. Nilufer Taner and Dr. Steven G. Younkin, Mayo Clinic, Jacksonville, FL using samples from the Mayo Clinic Study of Aging, the Mayo Clinic Alzheimer’s Disease Research Center, and the Mayo Clinic Brain Bank.

We would like to thank Sudeshna Das, Colin Magdamo, Roy Welsch, Stan Finkelstein, Ioanna Tzoulaki, Deborah Blacker and Lefkos Middleton for their insightful comments and suggestions, and Paul Murrell for his relentless help with grImport2 and gridSVG packages that enabled complete figure reproducibility within R.

## Funding

We acknowledge support from the NIA grant R01 AG058063 (SR, CH, PT, AS, MWA, BH and PKS), U54-CA225088 and a supplement CA225088-02S1 (SR, MWA, NTJ, KE, PKS), U24-DK116204 (NM, SAB and PKS) the CART fund (awarded to M.W.A), and by Harvard Catalyst Program for Faculty Development and Diversity Inclusion (PFDD) Faculty Fellowship (awarded to S.R.)

## Author contributions

SR, SB, KE and GZ collected transcriptional perturbation and drug toxicity data. PT and AS implemented the machine learning framework. CH and NM conducted drug target analysis. All authors contributed to figures and manuscript writing.

## Competing interests

PKS is a member of the SAB or Board of Directors of Applied Biomath, RareCyte Inc and Glencoe Software and has equity in these companies. In the last five years the Sorger lab has received research funding from Novartis and Merck. PKS declares that none of these relationships are directly or indirectly related to the content of this manuscript. BTH has stock in Novartis and Dewpoint. NTJ is an employee of H3 Biomedicine, a subsidiary of Eisai Inc. that develops therapies for Alzheimer’s. SR, PKS, MWA, and AS are inventors on a patent application for novel targets in neurodegenerative diseases.

---

**Supplementary Fig. 1:**
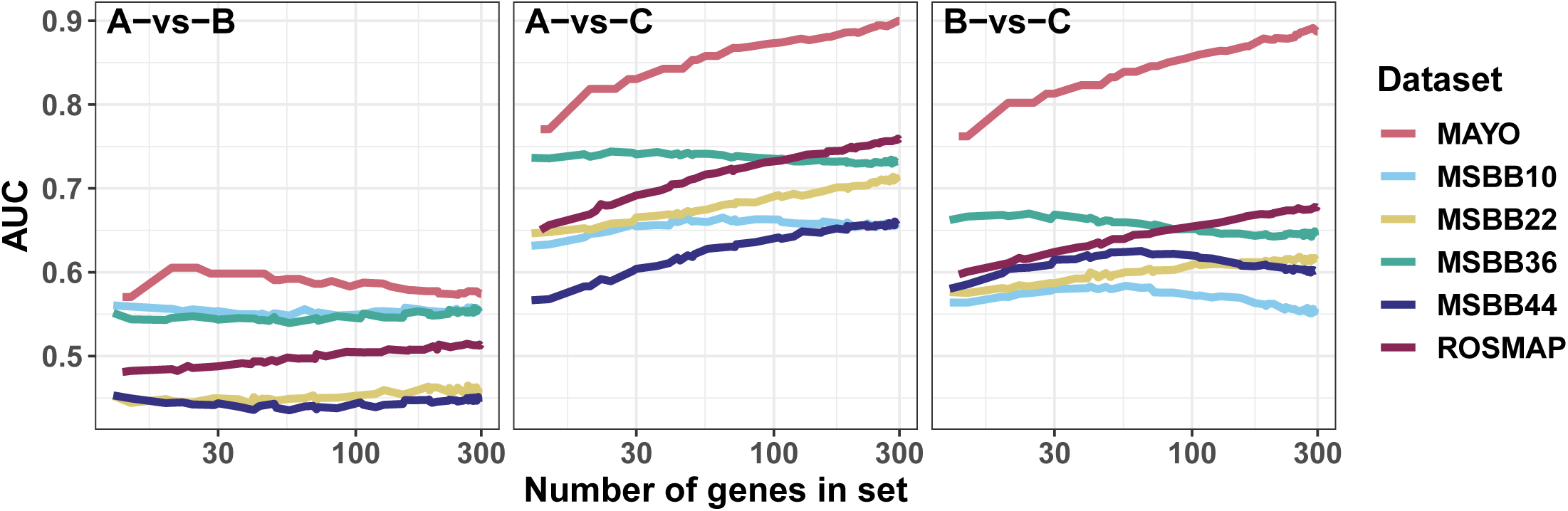
Performance of predictors trained on randomly-selected sets of genes plotted as a function of the set size. Performance was evaluated through leave-pair-out cross-validation and displayed as area under the ROC curve (AUC). The three panels correspond to binary classification tasks comparing early (A), intermediate (B) and late (C) disease stages. The color scheme, as introduced in Fig. 1b, denotes the dataset and brain region (specified as Brodmann Area) of samples used in each analysis.

**Supplementary Fig. 2:**
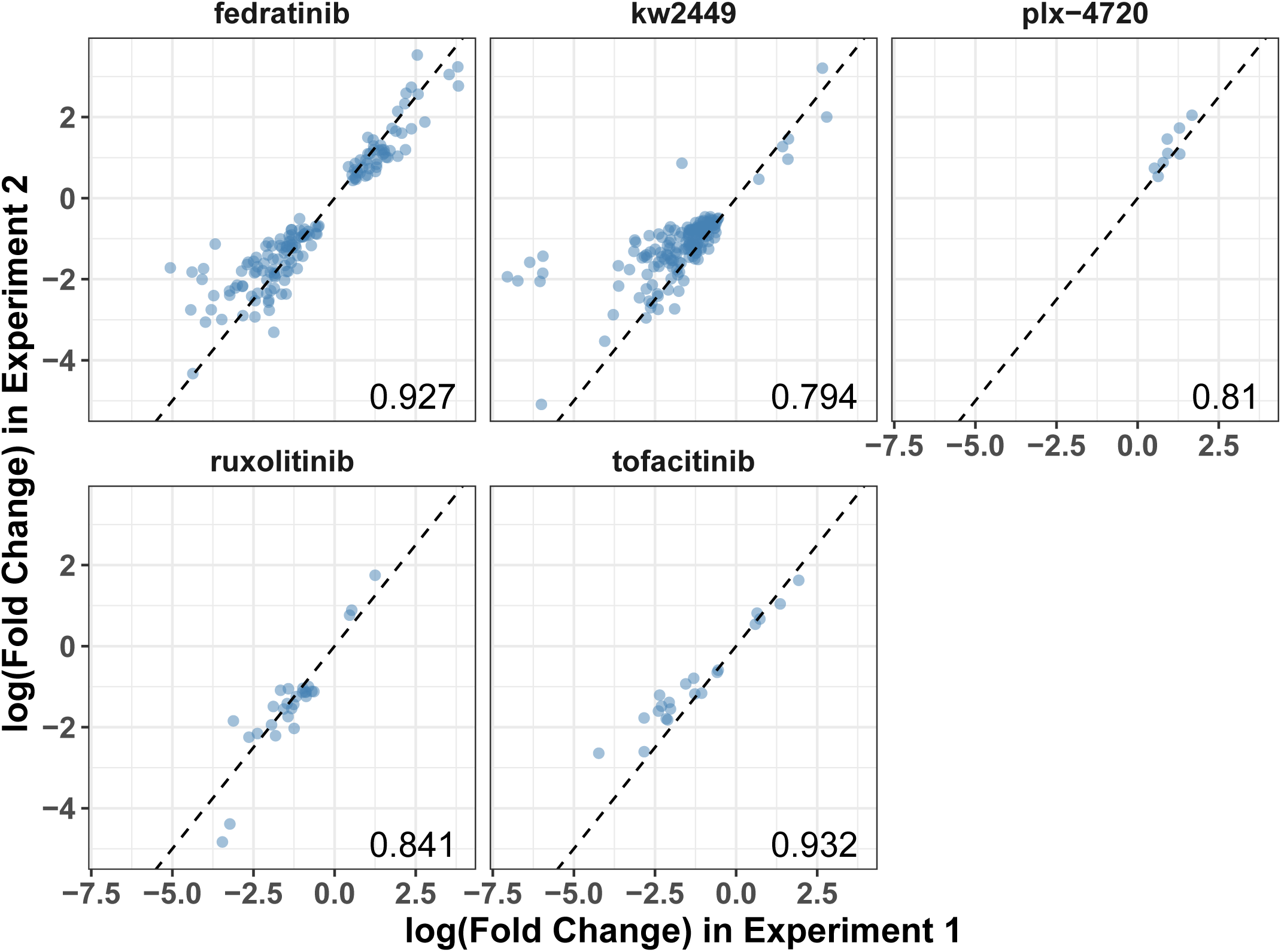
Concordance of treatment replicates across the two 3-DGE experiments. Shown are log-fold change values for all genes that were significantly (FDR ≤ 0.05) perturbed in both 3-DGE experiments. Spearman correlation between the two experiments is displayed in the bottom right corner of each panel.

**Supplementary Fig. 3:**
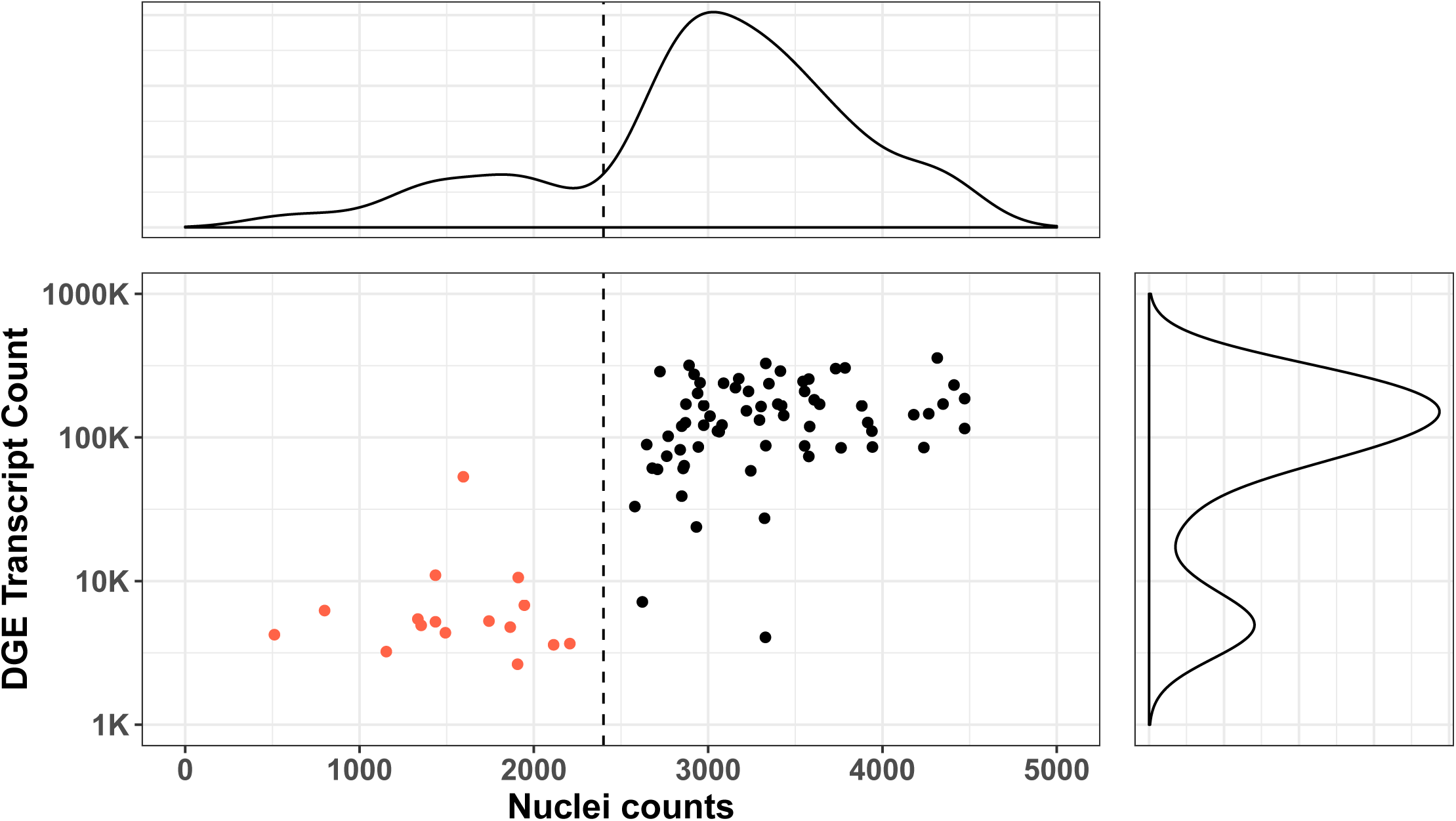
Assessment of compound toxicity. Nuclei counts estimated from microscopy images (x-axis) are plotted against mRNA abundance (y-axis). The mRNA abundance was computed as the total number of transcripts in the post-perturbational gene expression profile of the corresponding compound. Marginal distributions presented on the top and the right-hand side exhibit bi-modality, suggesting natural thresholds for determining compound neurotoxicity. A vertical dashed line is used to classify compounds into Toxic and Non-Toxic categories for Fig. 3.

**Supplementary Fig. 4:**
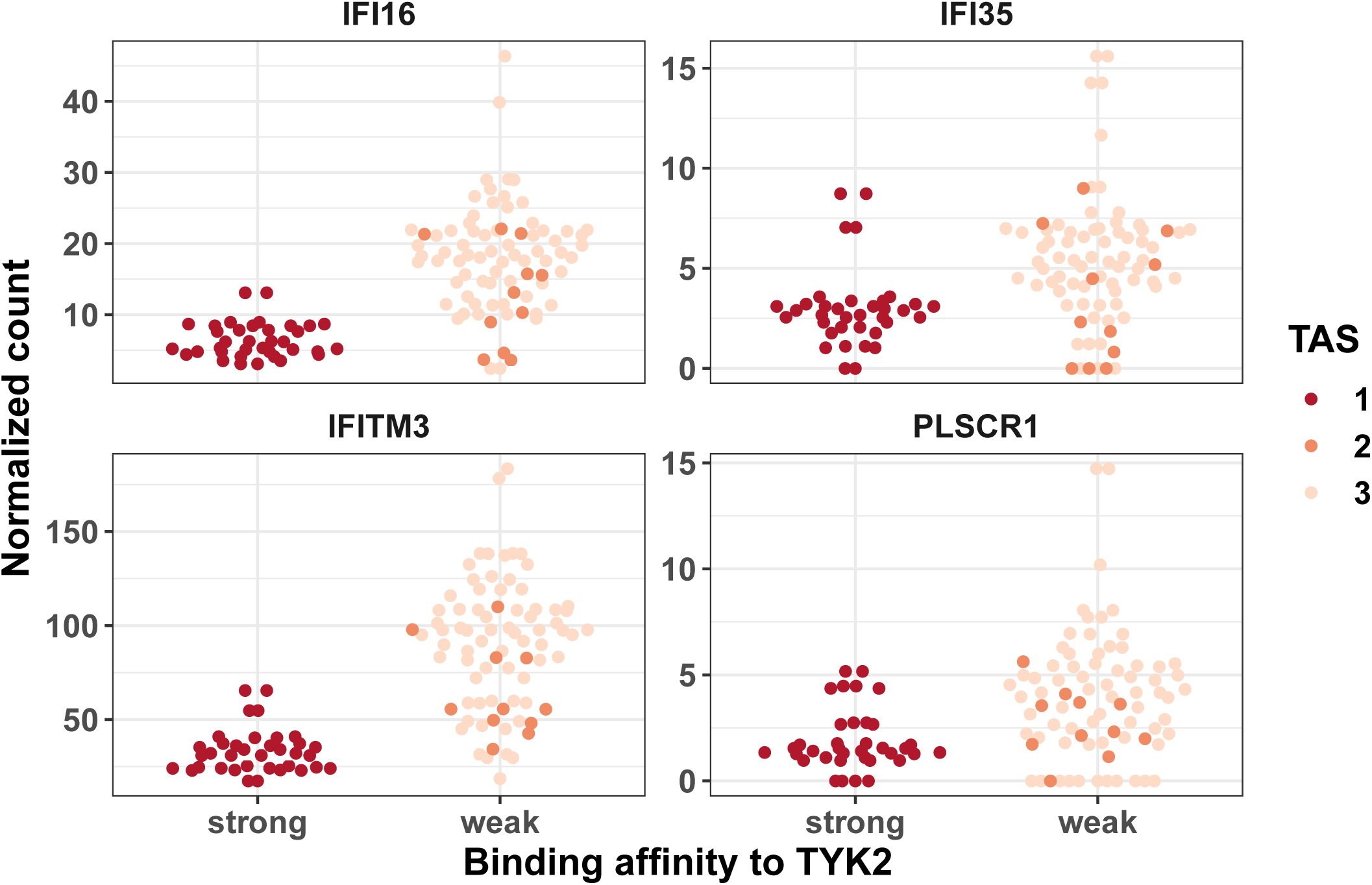
Raw expression values of selected interferon-stimulated genes. Each panel shows normalized transcript counts for a single interferon-stimulated gene (ISG). Individual points correspond to compounds that have a strong (TAS=1) or weak (TAS=2,3) binding to TYK2. Direct comparison of expression distributions (Wilcoxon Rank Sum test) between strong and weak binders was observed to be statistically significant (p ≤ 1e-4) for all four genes.

## Notes

https://synapse.org/DRIAD

https://github.com/labsyspharm/DRIAD

https://github.com/labsyspharm/DRIADrc

